# Structural insights into the specific recognition of H2A.Z-H2B dimer by the catalytic subunit of SRCAP chromatin remodeling complex

**DOI:** 10.64898/2025.12.30.697126

**Authors:** Wansen Tan, Bingyan Yuan, Jingjun Hong

## Abstract

Chromatin remodeling complexes (SRCAP-C) mediate the exchange of H2A.Z with H2A, leading to the incorporation of H2A.Z-H2B dimers into chromatin through a step-by-step manner. Here, we determined the crystal structure of the N-terminal region (531-560) of the catalytic subunit of SRCAP-C, termed SRCAP-Z domain, in complex with the engineered single-chain H2B-H2A.Z (scH2B-H2A.Z) at 2.53 Å resolution. In the crystal structure, two molecules of scH2B-H2A.Z and one molecule of SRCAP-Z form a sandwich-like architecture. Histones form typical histone fold and SRCAP-Z domain adopts a short 3_10_ helix. In details, Val551, Leu554, and Leu555 of the SRCAP-Z 3_10_ helix specifically recognize the αC helix of H2A.Z through hydrophobic interactions. The particular recognition has been validated *in vitro* by MBP-pulldown and isothermal titration calorimetry (ITC) experiments, and the MBP-pulldown assay results demonstrated that mutations in specific amino acids of SRCAP-Z attenuated its interaction with H2A.Z-H2B dimer. Finally, we demonstrated that SRCAP-Z recognizes H2A.Z-H2B dimer with a conservative structural basis and facilitates the nucleosome assembly by its chaperone activity.

**Highlights:**

Firstly, we determined the structure of the SRCAP-Z domain/scH2A.Z-H2B complex at 2.53 Å resolution.
Second, our results support a mechanism of H2A.Z-H2B/H2A-H2B exchange model in which YL1 enriches H2A.Z/H2B around nucleosomes, presenting them to SRCAP, which mediates the exchange of H2A.Z/H2B with H2A/H2B.
Third, SRCAP-Z recognition of H2A.Z-H2B dimer is structurally conserved.
Fourth, a distinct sandwich-like architecture: SRCAP-Z adopts a 3_10_ helix that inserts precisely between H2A.Z and H2B, forming a stable, unexpected structural module.

## Introduction

As the primary carrier of genetic information in eukaryotes, chromatin’s precise regulation of its structure and function is crucial for life activities. The dynamic changes of chromatin are closely related to various biological processes, including gene transcription (Giaimo et al., 2019, Altaf et al., 2010, Vlijm et al., 2015), DNA replication, DNA damage repair, and so on. Within the chromatin structure, nucleosomes serve as its fundamental building blocks, and the composition and modifications of nucleosomes significantly influence chromatin function (Luger et al., 1997). Chromatin remodeling complexes, which rely on ATPase for energy, are capable of sliding, exchanging, or even altering the composition of nucleosomes. Chromatin remodeling complexes are mainly classified into four major categories: ISWI (Barisic et al., 2019), CHD (Klement et al., 2014), SWI/SNF (Johnston et al., 1999, Mizuguchi et al., 2004), and INO80 (Hong et al., 2014). The INO80 family encompasses two types of complexes, INO80 and SWR1 (Morrison and Shen, 2009). The INO80 and SWR1 complexes share similar subunit compositions and highly conserved structural domains. Unlike the INO80 chromatin remodeling protein, the SWR1 complex cannot move nucleosomes. The SRCAP complex (SRCAP-C), p400/TIP60 complex, and INO80 complex belongs to the INO80. The SRCAP complex is a super-macromolecular chromatin remodeling complex composed of ten components: SRCAP, DMAP1, YL1, RUVBL1, RUVBL2, ACTL6A, ARP6, ACTIN, GAS41, and ZNHIT1 (Yu et al., 2024, Park et al., 2024). The distribution of H2A.Z in the genome exhibits specificity, primarily enriching in dynamic nucleosomes located at gene promoters and enhancer regions. This distribution pattern renders it an important marker for gene regulation. The formation of H2A.Z nucleosomes is dependent on the action of the chromatin remodeling complex SRCAP /SWR1, a process that requires ATP consumption and is facilitated by specific histone chaperones. The SRCAP complex recognizes and binds to the H2A.Z-H2B dimer through its specific subunits, and the structural basis for this process has gradually been elucidated. Studies have shown that the YL1 subunit of the SRCAP complex plays a pivotal role in H2A.Z recognition (Hong et al., 2025, Liang et al., 2016, Latrick et al., 2016).

The process by which the SRCAP complex mediates the integration of the H2A.Z-H2B dimer into chromatin involves complex molecular mechanisms (Wong et al., 2007). Studies have demonstrated that the exchange of the H2A.Z variant entails the removal of the H2A-H2B dimer from pre-assembled nucleosomes and its replacement with the H2A.Z-H2B dimer. A specific family of chromatin remodeling complexes (homologues of the yeast Swr1 complex) has been shown to be capable of performing this exchange. As a member of this family, SRCAP plays a crucial role in the integration process of H2A.Z. Research has found that the SRCAP complex facilitates the deposition of the histone variant H2A.Z in an ATP-dependent manner, which is essential for transcriptional regulation (Ruhl et al., 2006), chromatin accessibility, and neurodevelopmental processes (Johal et al., 2025).

Though the overall structure of the SRCAP complex was resolved by cryo-EM (Yu et al., 2024, Park et al., 2024), the structural basis or mechanism by which SRCAP exchanges of H2A with H2A.Z remain unclear. Our previous work found that the human SRCAP_501-607_ region may interact with human H2A.Z-H2B dimer(Hong et al., 2014), however, the structural basis is still unknown. Here, we clearly revealed how human SRCAP_531-560_ specifically recognizes the human H2A.Z-H2B dimer by a conserved structural basis or mechanism.

## Results

### Structure of the SRCAP and single chain H2B-H2A.Z complex

To investigate the structural basis for the recognition of H2A.Z by SRCAP subunits, X-ray crystallography was employed to determine the structure of the SRCAP_531-560_/scH2B-H2A.Z complex. Here we have determined the crystal structure of the SRCAP complex subunit, in complex with the scH2B-H2A.Z dimer at 2.53 Å resolution (PDB ID: 9UBD, Figure 1A-B and Table 1). In the crystal structure, two molecules (molecule 1 and molecule 2) of scH2B-H2A.Z and one molecule of SRCAP_531-560_ form a sandwich-like architecture (Figure 1B-C). The amino acid residues 531-560 of SRCAP, was termed as SRCAP-Z domain (Figure 1D). The SRCAP-Z domain residues 551-556, a 3_10_ helix, interact with the residues in the α3 and αC helices of H2A.Z. Typically, Val551, Leu554, and Leu555 of the SRCAP-Z specifically recognize the residues Ile90 in the α3 helix and Ile100 and Ile104 in the αC helix of H2A.Z through hydrophobic interactions, respectively (Supplementary Figure S1C).

**Figure 1.**
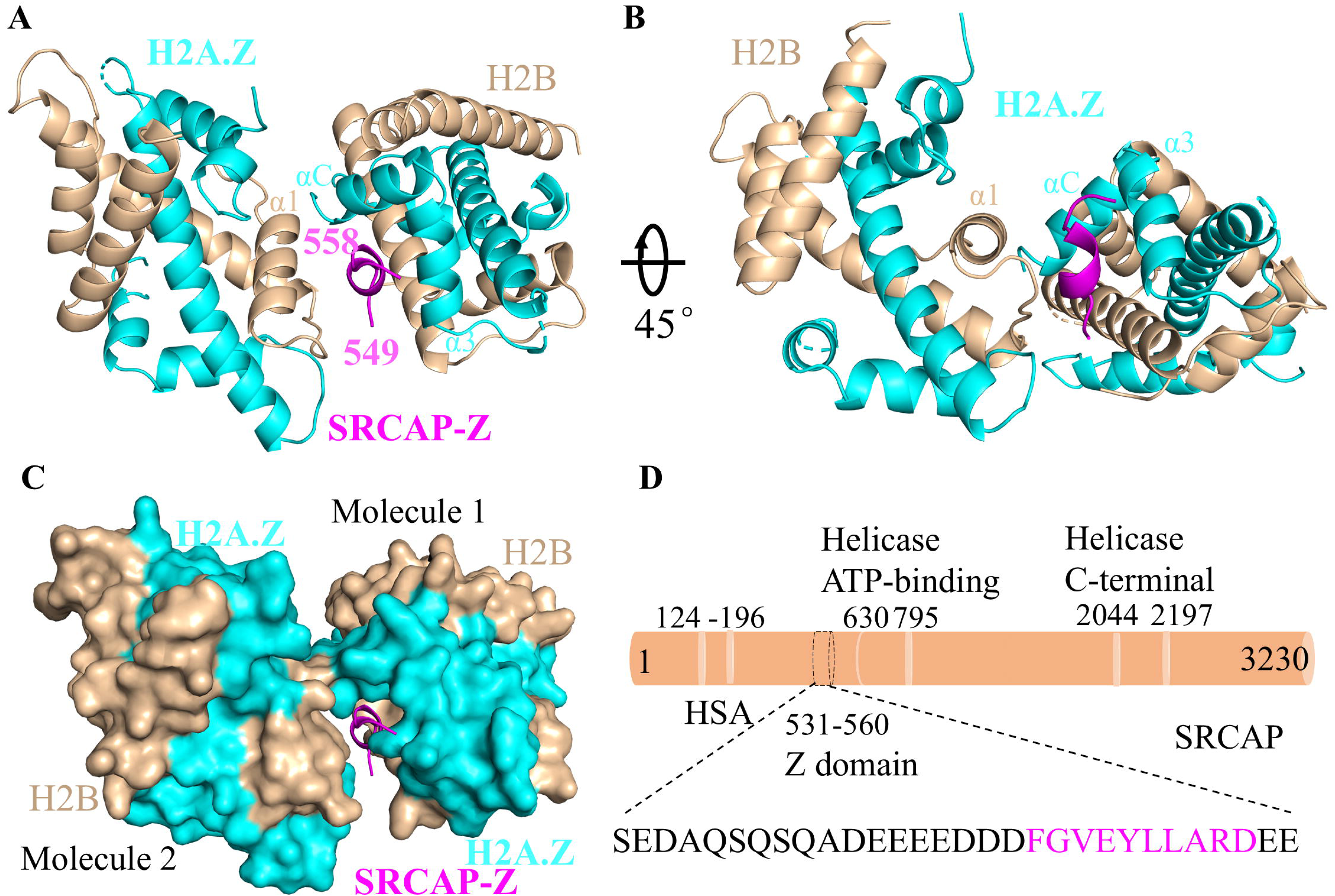
Structure of the SRCAP_531-560_/scH2B-H2A.Z complex. (A-B) Overall structure of the SRCAP_531-560_/scH2B-H2A.Z complex. Magenta: SRCAP-Z; Cyan: H2A.Z; Wheat: H2B; Figure B shows the overall structure after rotation. (C) Surface view of the SRCAP_531-560_/scH2B-H2A.Z complex, where SRCAP-Z is sandwiched between two scH2B-H2A.Z molecules to form a sandwich structure. (D) Illustration of the amino acid sequence in the region 531-560 of the SRCAP subunit. The magenta marked amino acid sequence represents the SRCAP-Z sequence in the complex structure.

**Table 1.** Data collection and refinement statistics. Optimized crystals (Supplementary Figure S1A) for data collection.

| *Human SRCAP-Z/scH2B-H2A.Z* |  |
| --- | --- |
| **Data collection** |  |
| Space group | C 1 2 1 |
| Cell dimensions |  |
| a,b,c(Å) | 77.33, 102.19, 67.37 |
| $\alpha,\beta,\gamma(^{\circ})$ | 90.00, 114.78, 90.00 |
| Molecules/asymmetric unit | 2 |
| Resolution (Å) | 35.10-2.53 |
| Rmerge<sup>a</sup> | 0.129 (0.645) |
| I/σI | 10.8 (2.6) |
| Completeness (%) | 92.7 (98.3) |
| Redundancy | 6.4 (6.2) |
| **Refinement** |  |
| Rwork | 0.252 |
| Rfree | 0.288 |
| Average B-factor (Å<sup>2</sup>) | 35.2 |
| Total number of atoms | 2685 |
| RMSD from ideality | 0.37 |
| Bond lengths (Å) | 0.51 |
| Bond angles (deg.) | 0.78 |
| Ramachandran analysis |  |
| Most-favored regions (%) | 94 |
| Additionally allowed regions (%) | 6 |

Note: scH2B-H2A.Z: single chain *Human* H2B-H2A.Z.

### Purification and identification of the MBP-SRCAP_521-569_/H2A.Z/H2B complex

To validate this interaction, wild-type proteins were expressed in *Escherichia coli* BL21(DE3). MBP-SRCAP_521-569_ (Figure 2A-B and Supplementary Figure S2A-C) and SUMO-SRCAP_521-569_ (Supplementary Figure S2D) were subsequently prepared by Ni-NTA, gel filtration chromatography. In parallel, H2A.Z/H2B was prepared (Supplementary Figure S3A-B), and gel filtration chromatography confirmed the formation of a complex between MBP-SRCAP_521-569_ and H2A.Z/H2B. MBP-SRCAP_521-569_ formed a complex with H2A.Z/H2B, as reflected by an earlier elution time compared to either component alone; in contrast, MBP alone did not form a detectable complex with H2A.Z/H2B (Figure 2C). It is demonstrated that MBP does not affect the interaction between SRCAP_521-569_ and H2A.Z/H2B. Dynamic light scattering (DLS) measurements showed an average particle size of 9.9 nm for the MBP-SRCAP_521-569_/H2A.Z/H2B complex (Supplementary Figure S3C-E). Isothermal titration calorimetry (ITC) measurements revealed a binding affinity of 256 nM for the interaction between SUMO-SRCAP_521-569_ and H2A.Z/H2B (Figure 2D).

**Figure 2.**
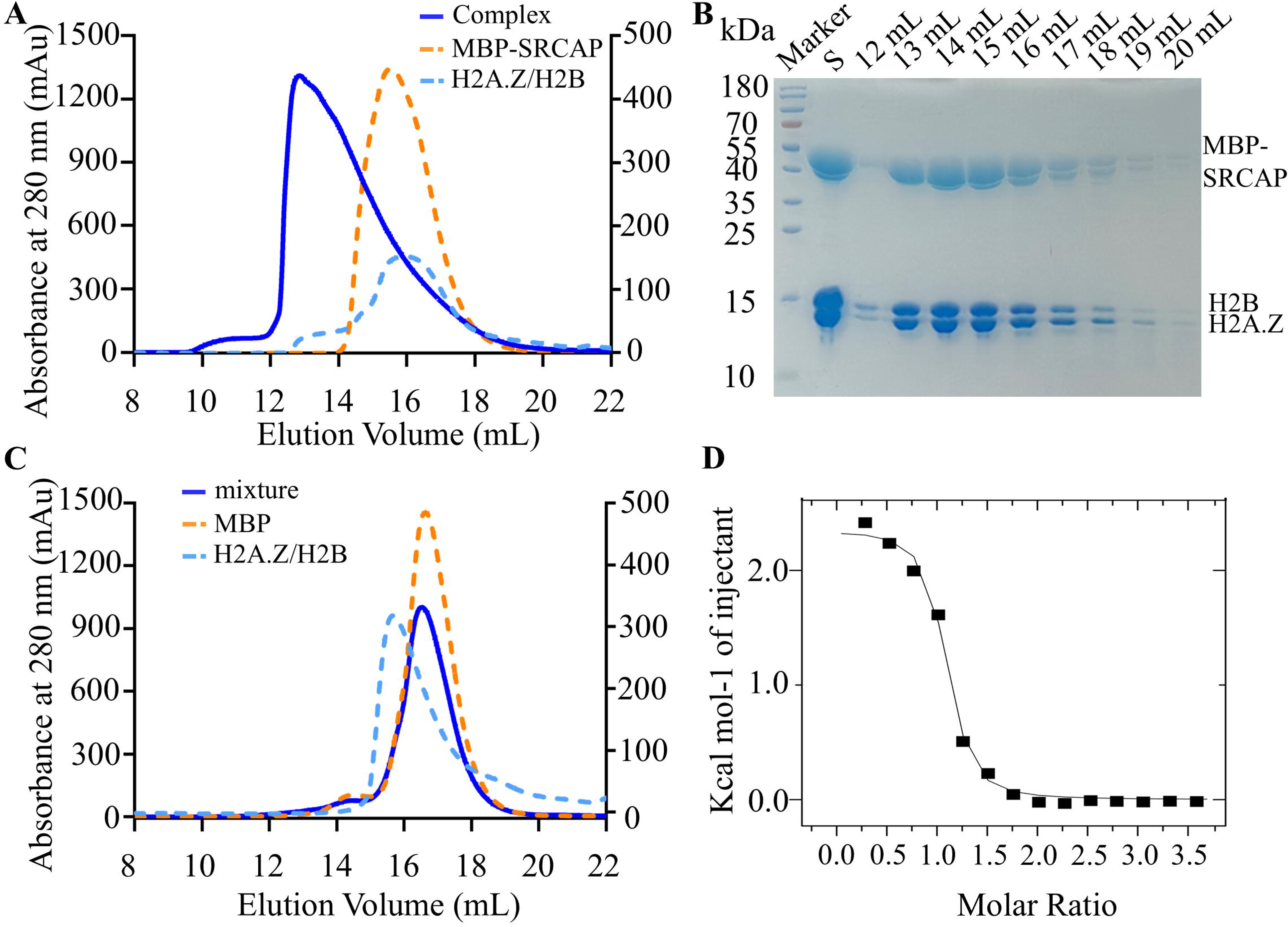
*In vitro* validation of the interaction between SRCAP_521-569_ and H2A.Z/H2B dimer (A) Gel flitration of the MBP-SRCAP_521-569_/H2A.Z/H2B complex. Blue represents the complex, cyan represents H2A.Z/H2B, orange represents MBP-SRCAP_521-569_, the peak position of the complex shifts forward. (B) SDS-PAGE of the MBP-SRCAP_521-569_/H2A.Z/H2B Complex. Marker, protein standard molecular weight; S, sample prior to gel filtration chromatography; 12-20 mL, elution volume. (C) Gel flitration of the MBP/H2A.Z/H2B mixture. (D) ITC profiles of the titration of the human SUMO-SRCAP_521-569_ with human H2A.Z/H2B. The binding affinities between SUMO-SRCAP_521-569_ and H2A.Z/H2B, as measured by ITC, are 256 nmol/L.

### Key residues in the interaction between scH2B-H2A.Z and SRCAP-Z

To identify the residues important for interactions between the H2A.Z/H2B and SRCAP, we generated a series of H2A.Z mutants targeting distinct sites, including single mutations Ile90Ala (I90A), Ile100Ala (I100A), Ile104Ala (I104A); double mutations such as I90A/I100A, I90A/I104A, I100A/I104A; Triple mutation such as I90A/I100A/I104A (Figure 3A).

**Figure 3.**
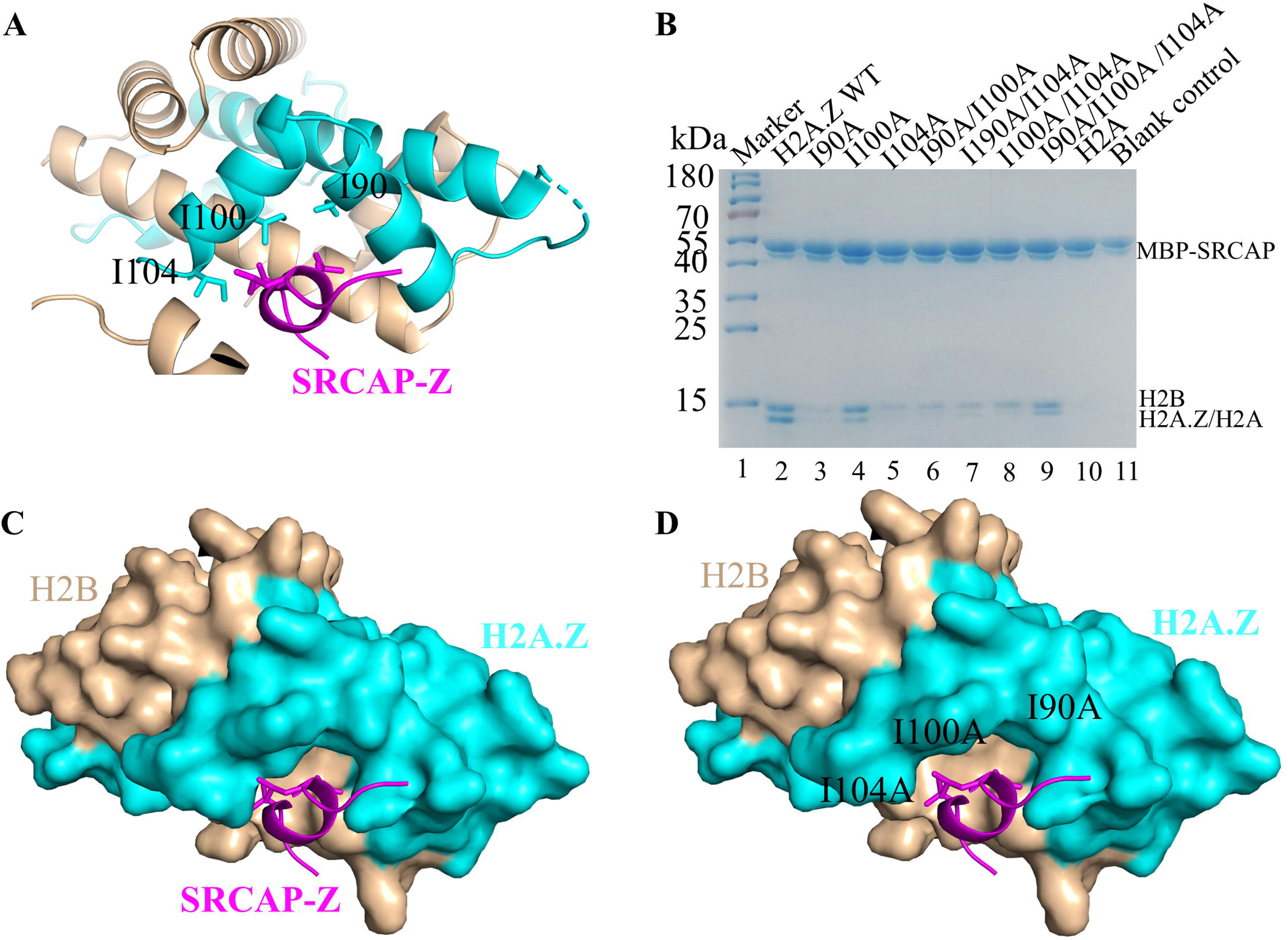
Specific interactions between the SRCAP-Z domain and scH2B-H2A.Z (A) Key amino acids in H2A.Z that maintain hydrophobic interactions with SRCAP-Z. (B) MBP pulldown of MBP-SRCAP_521-569_ and H2A.Z/H2B complexes with WT H2A.Z and the indicated H2A.Z mutants. Marker, protein standard molecular weight; H2A.Z WT, wild-type H2A.Z/H2B dimer; I90A, I100A, I104A, I90A/I100A, I90A/I104A, I100A/I104A, I90A/I100A/I104A, mutants of H2A.Z/H2B dimer; H2A, wild-type H2A/H2B dimer; Blank control, do not add H2A.Z/H2B dimer. See also Figure S4A. (C) Space position of SRCAP-Z and Wild-Type scH2B-H2A.Z. (D) Space position of SRCAP-Z and mutant scH2B-H2A.Z.

MBP-pulldown assays with H2A.Z/H2B and MBP-SRCAP_521-569_ showed that each single mutant and the combinatorial mutants weakened binding (Figure 3B, lanes 3-9 and Supplementary Figure S4A). Compared with the wild type, the single mutant I100A did not significantly reduce binding to MBP-SRCAP_521-569_. I100 is located between I90 and I104 (Figure 3A); mutation of I100 to Ala preserves the hydrophobic interactions mediated by I90 and I104 with SRCAP-Z, thereby maintaining the binding of SRCAP-Z to the I100A mutant. The triple mutant I90A/I100A/I104A also did not exhibit a significant reduction in binding to SRCAP-Z. We propose that when I90, I100, and I104 are all mutated to Ala, steric hindrance will decrease, allowing SRCAP-Z to approach the αC and α3 helices of H2A.Z more closely (Figure 3C-D). These results are fully consistent with the crystal structure. We conclude that Ile90, Ile100, and Ile104 are the key amino acids that maintain the hydrophobic interaction between H2A.Z and SRCAP-Z.

### Residues important for interactions between the SRCAP-Z domain and scH2B-H2A.Z

To further identify key residues in the interaction between the SRCAP-Z domain and H2A.Z/H2B, we generated a series of MBP-SRCAP_521-569_ mutants targeting distinct sites, including single mutations Val551Ala (V551A), Leu554Ala (L554A), Leu555Ala (L555A), double mutations V551A/L554A, V551A/L555A, L554A/L555A, and a triple mutation V551A/L554A/L555A (Figure 4A).

**Figure 4.**
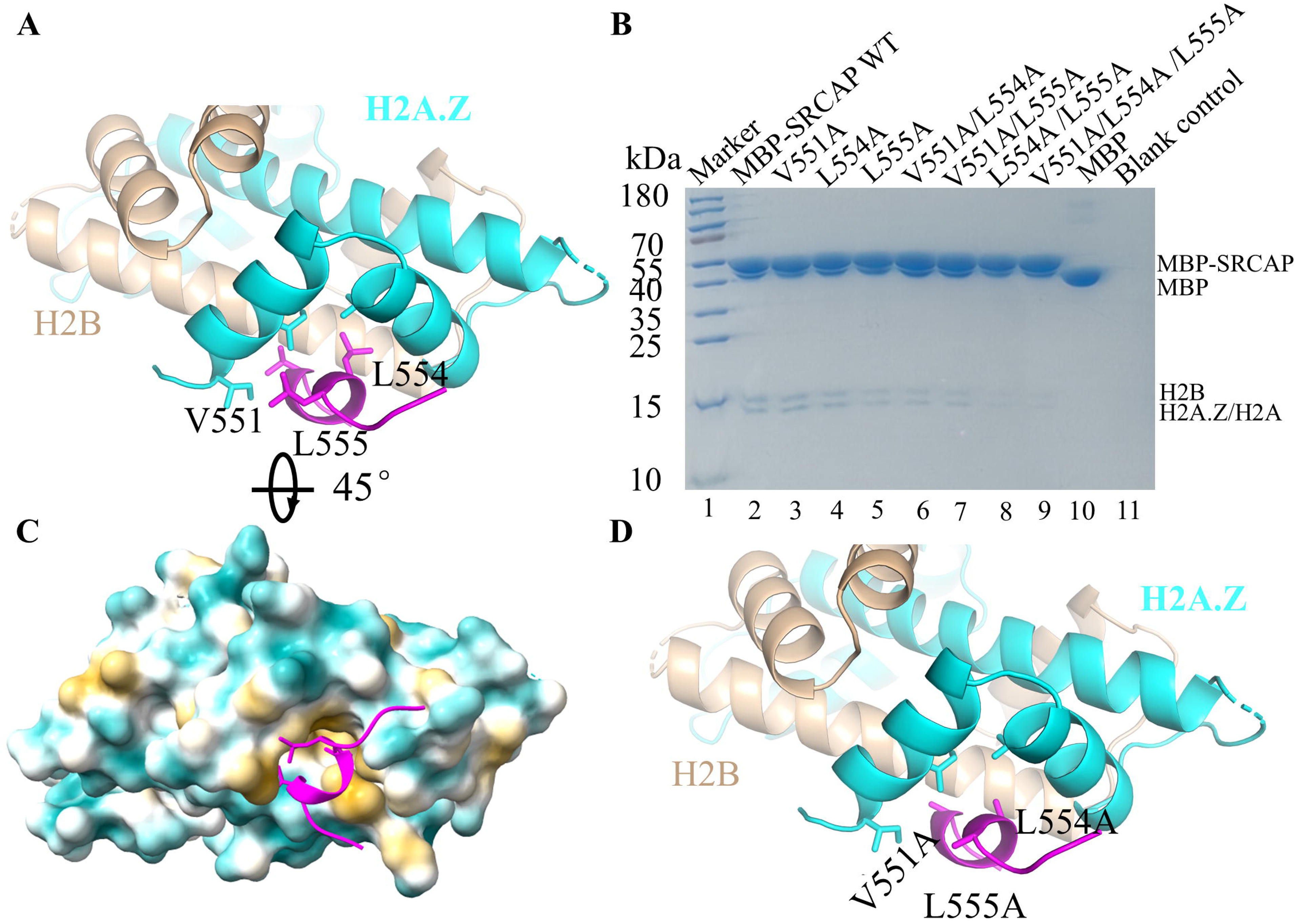
Residues important for interactions between the SRCAP-Z domain and scH2B-H2A.Z (A) Key amino acids in SRCAP-Z that maintain hydrophobic interactions with H2A.Z (B) MBP-pulldown of MBP-SRCAP_521-569_ (mutants) and H2A.Z/H2B dimer. Marker, protein standard molecular weight; MBP-SRCAP WT, wild-type MBP-SRCAP_521-569_; V551A, L554A, L555A, V551A/L554A, V551A/L555A, L554A/L555A, V551A/L554A/L555A, mutants of MBP-SRCAP_521-569_; MBP, wild-type MBP; Blank control, do not add MBP-SRCAP_521-569_. See also Figure S4B. (C) SRCAP-Z is located within the hydrophobic region of H2A.Z. Cyan: Hydrophilic residues; Yellow: Hydrophobic residues. (D) SRCAP-Z key amino acid residue mutated to Ala.

MBP-pulldown assays demonstrated that binding to H2A.Z/H2B was impaired by the V551A, L554A, and L555A mutations. The double mutants V551A/L554A, V551A/L555A, and L554A/L555A exhibited substantially reduced binding to H2A.Z/H2B (Figure 4B, lanes 3-9 and Supplementary Figure S4B). Notably, the triple mutant V551A/L554A/L555A nearly completely lost binding. These results indicate that residues V551, L554, and L555 collectively contribute to the hydrophobic interaction with the H2A.Z-H2B dimer (Figure 4C). We conclude that Val551, Leu554, and Leu555 are required and important for the specific recognition of H2A.Z/H2B by the SRCAP-Z domain (Figure 4D).

### SRCAP-Z recognition of H2A.Z-H2B dimer is structurally conserved

To further understand the structural basis of SRCAP recognition of H2A.Z/H2B. We analyzed the complexes of H2A.Z/H2B bound to Swr1 (PDB ID: 4M6B), H2A.Z/H2B bound to Anp32e (PDB ID: 4NFT), and H2A.Z/H2B bound to SRCAP (PDB ID: 9UBD), respectively. The H2A.Z derived from yeast exhibits strong sequence conservation with its human, and H2A.Z is highly similar in sequence to H2A. The Swr1 subunit from *Saccharomyces cerevisiae* specifically recognizes H2A.Z/H2B using the intrinsically disordered Swr1-Z domain (Figure 5A). Similarly, *Homo sapiens* Anp32e also specifically recognizes the αC helix of H2A.Z (Figure 5B). Our determined complex structure reveals that SRCAP-Z specifically recognizes H2A.Z (Figure 5C). Interestingly, the sequences of swr1 and SRCAP/Anp32e that recognize H2A.Z show low sequence homology (Figure 5D and Supplementary Figure S5B), meaning that Swr1-Z and SRCAP-Z sequences are not similar. Despite different evolutionary origins, Swr1 from *Saccharomyces cerevisiae* and SRCAP/Anp32e from *Homo sapiens* use similar structures to recognize H2A.Z/H2B (Figure 5A-B). This indicates that Swr1 and SRCAP/Anp32e maintain structural conservation throughout evolution (Table 2).

**Figure 5.**
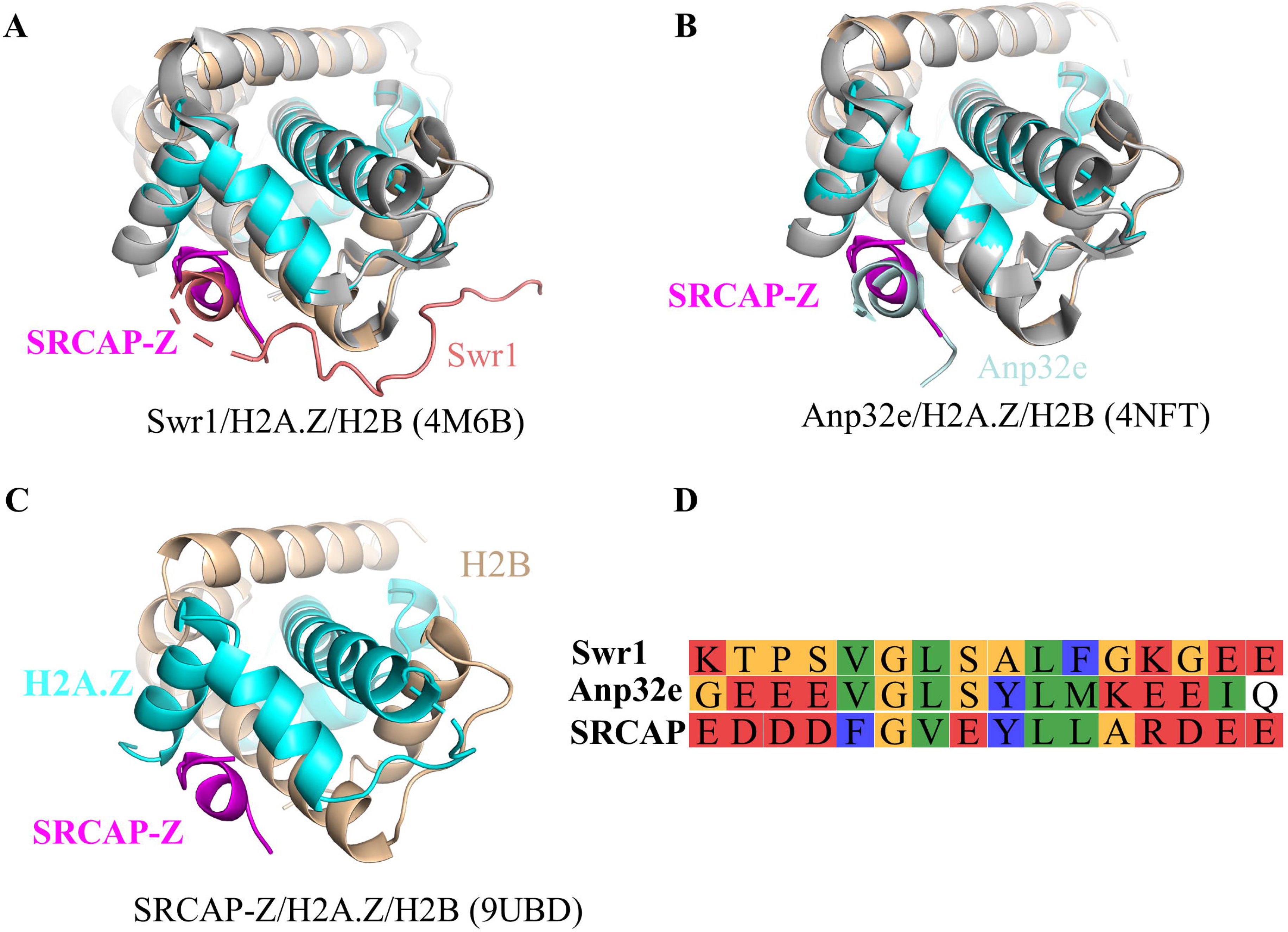
SRCAP-Z recognizes H2A.Z-H2B with structural conservation (A) SRCAP-Z/scH2B-H2A.Z complex structure. (B) Alignment of SRCAP-Z and Swr 1 structure. (C) Alignment of SRCAP-Z and Anp32e structure. (D) Amino acid residue comparison of the H2A.Z/H2B interacting regions among Swr1, Anp32e, and SRCAP.

**Table 2.** Structural basis for histone chaperone recognition of H2A.Z/H2B. Structural statistics of the nucleosome remodeling complex subunits and the H2A.Z/H2B complex.

|  | Experimental<br>method | resolution | Histone | Organism | PDB<br>ID | Reference | Conclusion |
| --- | --- | --- | --- | --- | --- | --- | --- |
| Swr1-Z | X-ray | 1.78 | H2B.1 and<br>H2A.Z | *Saccharomyces*<br>*cerevisiae*<br>S288C | 4M6B | Hong et<br>al., 2014 | The Swr1-Z domain<br>can deliver the<br>H2A.Z-H2B<br>dimer to the<br>DNA-(H3-H4)<sub>2</sub><br>tetrasome. |
| Anp32e-Z | X-ray | 2.61 | H2B type 2-E<br>and H2A.Z | *Homo sapiens* | 4NFT | Mao et<br>al., 2014 | Anp32e specifically<br>interacts with the<br>extended H2A.Z αC<br>helix. |
| YL1 | X-ray | 1.89 | scH2B-H2A.Z | *Drosophila*<br>*melanogaster* | 5CHL | Liang et<br>al., 2016 | The dYL1-Z domain<br>adopts a new whip-like<br>structure that wraps<br>over H2A.Z-H2B. |
| YL1 | X-ray | 2.70 | H2B TYPE<br>1-J and<br>H2A.Z | *Homo sapiens* | 5FUG | Latrick et<br>al., 2016 | YL1 binding, similarly<br>to Anp32e binding,<br>triggers an extension of<br>the H2A.Z αC helix. |
| YL1-Z | X-ray | 2.01 | scH2B-H2A.Z | *Homo sapiens* | 9INC | Hong et<br>al., 2025 | YL1-Z employ a<br>conserved structural<br>mechanism for<br>recognizing<br>H2A.Z-H2B dimer. |
| SRCAP-Z | X-ray | 2.53 | scH2B-H2A.Z | *Homo sapiens* | 9UBD | - | SRCAP-Z recognizes<br>H2A.Z-H2B dimer<br>with conservative<br>domain specificity. |

## Molecular mechanism of H2A.Z-H2B/H2A-H2B exchange in the nucleosome

The YL1-Z/scH2B-H2A.Z (PDB ID: 9INC)(Hong et al., 2025) and SRCAP-Z/scH2B-H2A.Z (PDB ID: 9UBD) complex structure from Professor Jingjun Hong’s group. Both YL1-Z(Hong et al., 2025, Liang et al., 2016, Latrick et al., 2016) and SRCAP-Z specifically recognize H2A.Z, SRCAP-Z and YL1-Z both play important roles in the exchange process between H2A.Z/H2B and H2A/H2B. To examine the functional role of the SRCAP_521-569_ domain, we performed *in vitro* reconstitution assays. Specifically, H2A.Z/H2B was incubated with MBP-SRCAP_521-569_ to form MBP-SRCAP_521-569_/H2A.Z/H2B trimer, which was then mixed with DNA-(H3.1-H4^A15C^)_2_ tetrasome (1:1 ratio) to assemble nucleosomes (Supplementary Figure S6A). After digestion with MNase enzyme, the tetrasome band was gone (disappeared) because it was more loose than NCP, while the NCP band remained, indicating that the NCP is more resistant to MNase mediated digestion (Supplementary Figure S6B), It has been confirmed that H2A.Z/H2B and H3.1/H4^A15C^ successful assembly into nucleosomes in the presence of SRCAP (Supplementary Figure S6A-B), it indicates that SRCAP can help H2A.Z/H2B form nucleosomes with H3/H4.

Based on these observations, when other subunits of the SRCAP complex are not considered, we propose a possible mechanistic model: The YL1 subunit of the SRCAP complex recognizes H2A.Z (Figure 6A). Subsequently, conformational changes bring H2A.Z/H2B into close proximity with the SRCAP subunit, H2A.Z/H2B/YL1/SRCAP, and other subunits (not shown here) and form a whole complex (Figure 6B). The complex approaches and engages the nucleosome (Figure 6C). Upon binding, the H2A/H2B dimer near super helical location 2 (SHL2) becomes partially detached from nucleosome DNA, adopting a partially engaged state(Park et al., 2024). H2A/H2B is exchanged for H2A.Z/H2B, resulting in the eviction of canonical H2A/H2B and the incorporation of H2A.Z/H2B into the nucleosome (Figure 6D). After the exchange is complete, the SRCAP complex dissociates, leaving behind a nucleosome containing H2A.Z/H2B (Figure 6E).

**Figure 6.**
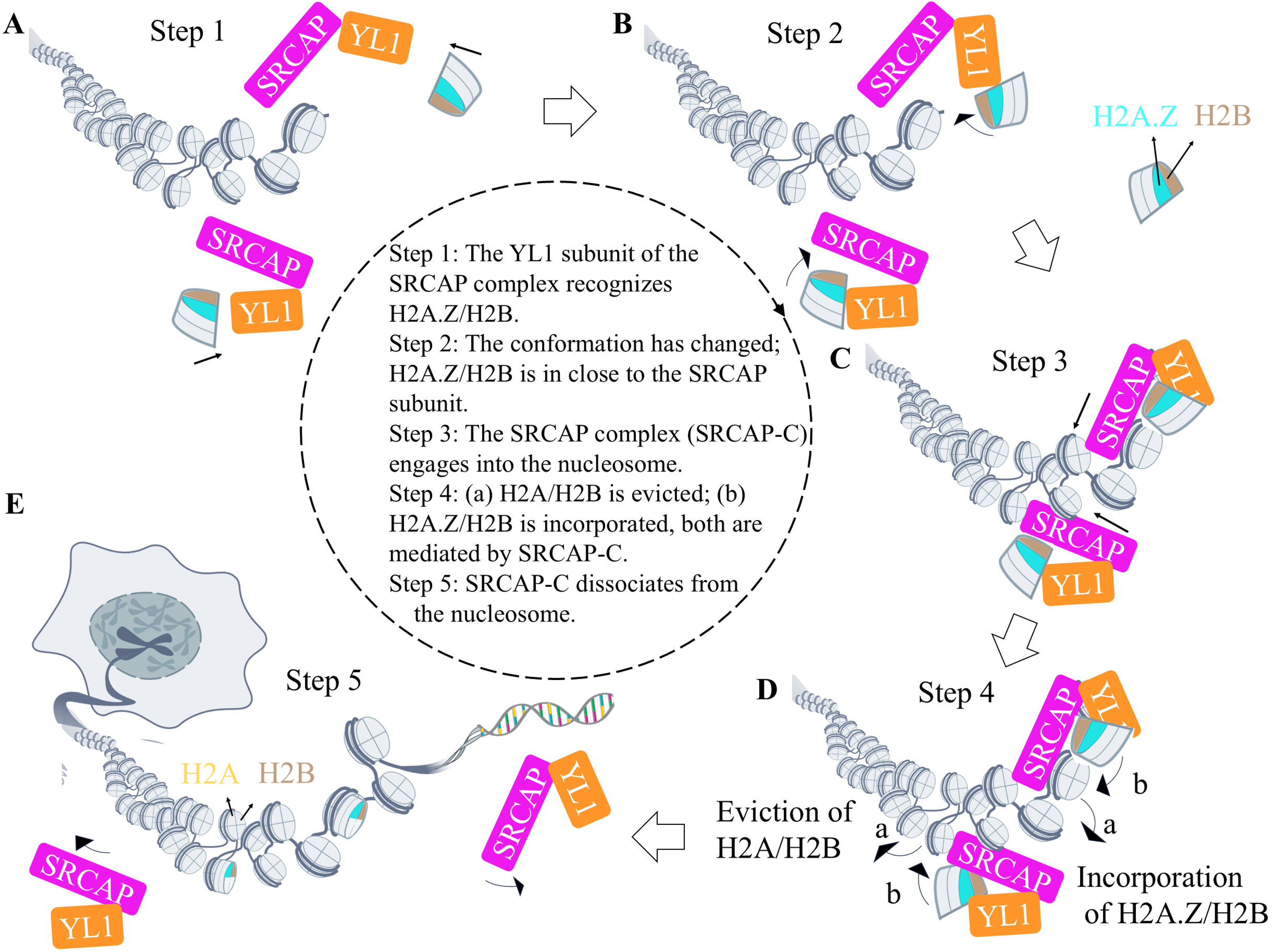
A proposed model for (H2A/H2B)/(H2A.Z/H2B) exchange mechanism by the SRCAP and YL1 (A) The YL1 subunit of the SRCAP complex recognizes H2A.Z/H2B, H2A.Z/H2B is enriched around YL1 (step1). (B) The conformation has changed, H2A.Z/H2B/YL1, SRCAP and other subunits (not shown here) and forms a whole complex (step2). (C) The SRCAP complex moves closer to the nucleosome (step3). (D) The H2A/H2B are evicted, H2A.Z/H2B is exchanged with H2A/H2B, both are mediated by SRCAP-C (step4). (E) The SRCAP complex dissociates from the nucleosome (step5). For simplicity, other subunits of the SRCAP remodeling complex were not taken into consideration.

## Discussion

In eukaryotic cells, dynamic chromatin regulation is achieved through multiple interconnected mechanisms, including DNA methylation, histone post-translational modifications, and the coordinated actions of histone chaperones(Venkatesh and Workman, 2015, Straube et al., 2010). The SRCAP complex functions as a key transcriptional regulator; Loss of SRCAP influences the timing of developmental gene expression(Johal et al., 2025).

In this study, we determined the crystal structure of the SRCAP-Z domain in complex with the scH2B-H2A.Z dimer, revealing the molecular basis for its specific recognition of H2A.Z. Through mutational analysis and MBP-pulldown assays, we identified critical residues within SRCAP-Z that mediate its interaction with the H2A.Z/H2B dimer. Interestingly, the I100A mutation only slightly weakens binding to SRCAP-Z, likely due to compensatory interactions involving I90 and I104. When all three residues (I90, I100, I104) are mutated to Ala, binding is not completely abolished, possibly due to reduced steric hindrance. Through structural comparisons, we found that Swr1-Z, Anp32e-Z, and SRCAP-Z responsible for recognizing the H2A.Z/H2B dimer exhibit structural conservation across evolution. In the interaction between SRCAP-Z and scH2B-H2A.Z, electrostatic interactions are not prominent (Supplementary Figure S7A-C). It is possible that regions outside SRCAP-Z engage in electrostatic interactions with scH2B-H2A.Z to stabilize the structure.

The SRCAP subunit within the SRCAP complex may be sterically hindered, limiting its accessibility to exogenous H2A.Z/H2B. In contrast, YL1 exhibits nanomolar affinity for H2A.Z/H2B (Hong et al., 2025), while SRCAP shows only micromolar affinity (Figure 2D). YL1 can readily sense surrounding H2A.Z/H2B with high sensitivity, making it more adept at recognizing H2A.Z/H2B in the external environment. We propose a possible model in which the YL1 subunit of the SRCAP complex first recognizes H2A.Z/H2B, before H2A.Z/H2B is recognized by YL1, it may be bound by histone chaperones such as Anp32e(Mao et al., 2014, Obri et al., 2014) and others. This recognition then leads to the activation of the H2A/H2A.Z exchange mechanism, followed by the SRCAP complex (SRCAP-C) engages into the nucleosome and the completion of the step-by-step exchange reaction (Figure 6).

It remains unclear whether SRCAP first recognizes H2A.Z before engages into the nucleosome or engages in the nucleosome first before interacting with H2A.Z. Another recognition and engagement mechanism may be the SRCAP complex first associating with the nucleosome, thereby preparing for the replacement of the H2A/H2B dimer.

Upon binding, the H2A/H2B dimer near super helical location 2 (SHL2) becomes partially detached from nucleosome DNA, adopting a partially engaged state(Park et al., 2024). Next step YL1 employs its N-terminal Z domain (YL1-Z) to recruitment the H2A.Z/H2B dimer. YL1 then delivers the H2A.Z/H2B dimer to the vicinity of the nucleosome, H2A.Z/H2B is enriched, YL1 presents it to the SRCAP. H2A.Z/H2B and H2A/H2B exchange mediated by SRCAP-C. After successful exchange, the SRCAP-C subsequently disengages from the nucleosome, completing one cycle of H2A.Z/H2B deposition, to further complete chromatin remodeling.

However, the exchange of H2A/H2B for H2A.Z/H2B and its associated chromatin remodeling is a complex process. For example, H2A.Z nucleosomes influence global chromatin architecture in a tail-dependent manner, which can be modulated by introducing the tail peptide into live cells(Imre et al., 2024). The involvement of the SRCAP remodeling complex in the complete chromatin remodeling requires further investigation.

## Experimental Procedures

### Cloning, expression, and purification of recombinant proteins

The DNA fragment encoding residues 521-569 of human SRCAP (*Homo sapiens* SRCAP Uniprot ID: Q6ZRS2) were amplified from the plasmid harboring the coding sequence of SRCAP and cloned into pET vectors harboring either an N-terminal MBP or SUMO tag. Site-directed mutagenesis was performed to generate mutant constructs. Recombinant plasmids were transformed into *Escherichia coli* BL21(DE3) cells, and protein expression was induced at low temperature to promote soluble expression. Proteins were purified sequentially using Ni-NTA agarose (Qiagen) and HiLoad^TM^ 16/600 Superdex^TM^ 200 pg gel filtration chromatography (GE Healthcare). Histone expression, purification, and assembly, refer to Professor Jingjun Hong’s article(Chittori et al., 2018).

The SRCAP_521-569_ amino acid sequence was: SASEESESEESEDAQSQSQADEEEEDDDFGVEYLLARDEEQSEADAGSG

### Crystallization, diffraction data collection, and structure determination

Peptide SRCAP_531-560_ was synthesized by GL Biochem (Shanghai) Ltd. SRCAP_531-560_ was mixed with purified scH2B-H2A.Z to form a stable complex. Crystals were grown at 16 °C using the sitting drop method. X-ray diffraction data were collected at the Shanghai Synchrotron Radiation Facility (SSRF) and processed using the automated data processing software. The structure was solved using the molecular replacement method.

The peptide SRCAP_531-560_ amino acid sequence was:

### SEDAQSQSQADEEEEDDDFGVEYLLARDEE

The single-chain H2B-H2A.Z (scH2B-H2A.Z) amino acid sequence used in this study was:

HMRKESYSIYVYKVLKQVHPDTGISSKAMGIMNSFVNDIFERIAGEASRLAHYNKRSTITSREIQTAVRLLLPGELAKHAVSEGTKAVTKYTSASSAVSRSQRAGLQFPVGRIHRHLKSRTTSHGRVGATAAVYSAAILEYLTAEVLELAGNASKDLKVKRITPRHLQLAIRGDEELDSLIKATIAGGGVIPHI*

### MBP-pulldown assays

MBP-SRCAP_521-569_ was conjugated to MBP besds *in vitro*. After incubation at 4°C, unbound proteins were washed away. H2A.Z/H2B was added for binding, followed by elution of the sample using elution buffer (0.2 mol/L NaCl, 7 mol/L Urea, 10 mmol/L Maltose) for subsequent SDS-PAGE analysis.

### Isothermal titration calorimetry (ITC) and Dynamic light scattering (DLS)

ITC experiments were performed using a MicroCal™ ITC200 instrument. Data were analyzed with ITC200 Analysis software to determine binding affinities and thermodynamic parameters. For DLS experiment: Prepare complexes of MBP-SRCAP with H2A.Z/H2B for particle size analysis and generate graphs using GraphPad.

### SRCAP assembly function experiment

Prepare nucleosomes by adding MBP-SRCAP_521-569_/H2A.Z/H2B dropwise to DNA/(H3.1/H4^A15C^)_2_ tetrasomes. Separate by DEAE chromatography, collect samples at different elution positions, digest with MNase, and performing native PAGE.

## Supplementary Materials

The following is the Supplementary data to this article:

## Accession code

Coordinates and structure factors have been deposited in the Protein Data Bank under accession codes PDB: 9UBD.

## Author Contributions

Prof. Jingjun Hong collected the financial funding, prepared the samples and optimization, conception and supervised the whole procedure of research, and paper writing. Wansen Tan (WT) performed crystal screening with the help of robots, WT and Prof. Hong performed crystal optimization. Prof. Hong, WT solved the SRCAP-Z/scH2B-H2A.Z structure. Bingyan Yuan (BY) and WT performed protein purification. BY performed pulldown assays, and ITC experiments. WT and Prof. Hong wrote the draft, and Prof. Hong revised the paper.

## Acknowledgments

This research was financially supported by the National Natural Science Foundation of China (grant 31970669) and the Research start-up funds of Anhui University (S020318006/067). We thank Congyu Liu and Dr. Tengchuan Jin (USTC) for their help with crystal data collection. We thank Dr. Jiansheng Jiang helped to build the scH2B-H2A.Z model using AlphaFold 2. We thank the staff of Beamline 02U1 at SSRF for assistance in data collection. We thank the core facility of School of Life Sciences and Medical Engineering, Anhui University.

