## Supplementary Data for "Structural insights into the specific recognition of H2A.Z-H2B dimer by the catalytic subunit of SRCAP chromatin remodeling complex"

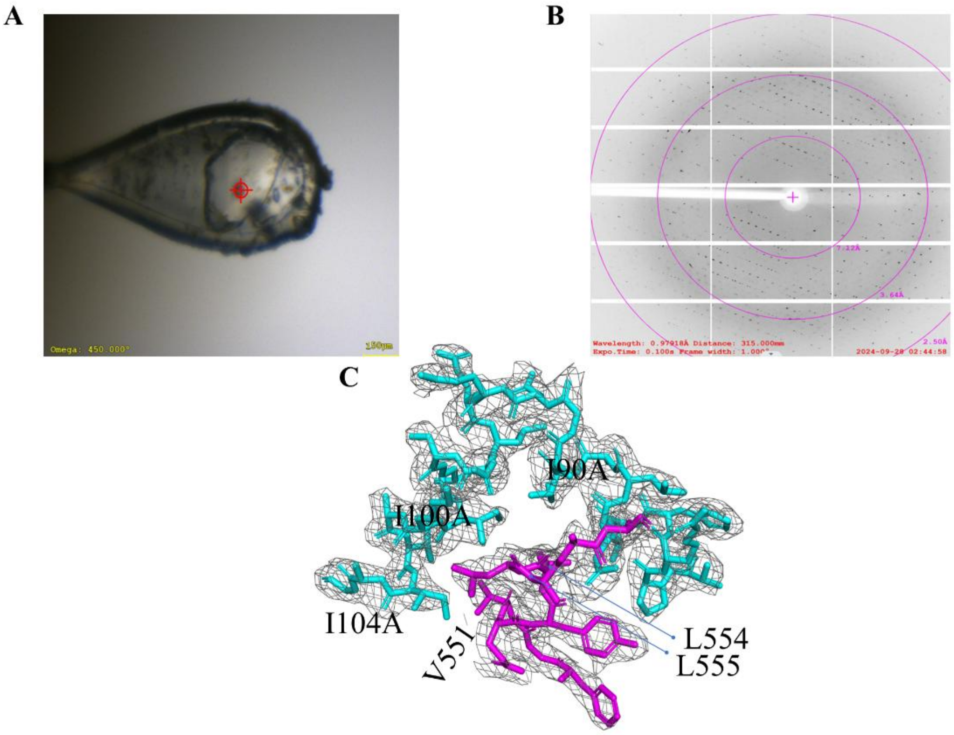


**Figure S1. Crystal data collection and density maps.**

A) Optimized SRCAP-Z domain/scH2B-H2A.Z complex crystals.

B) Crystal diffraction pattern of SRCAP-Z domain/scH2B-H2A.Z complex.

C) The omit map of SRCAP-Z and H2A.Z interaction positions. Cyan: H2A.Z; Magenta: SRCAP-Z.

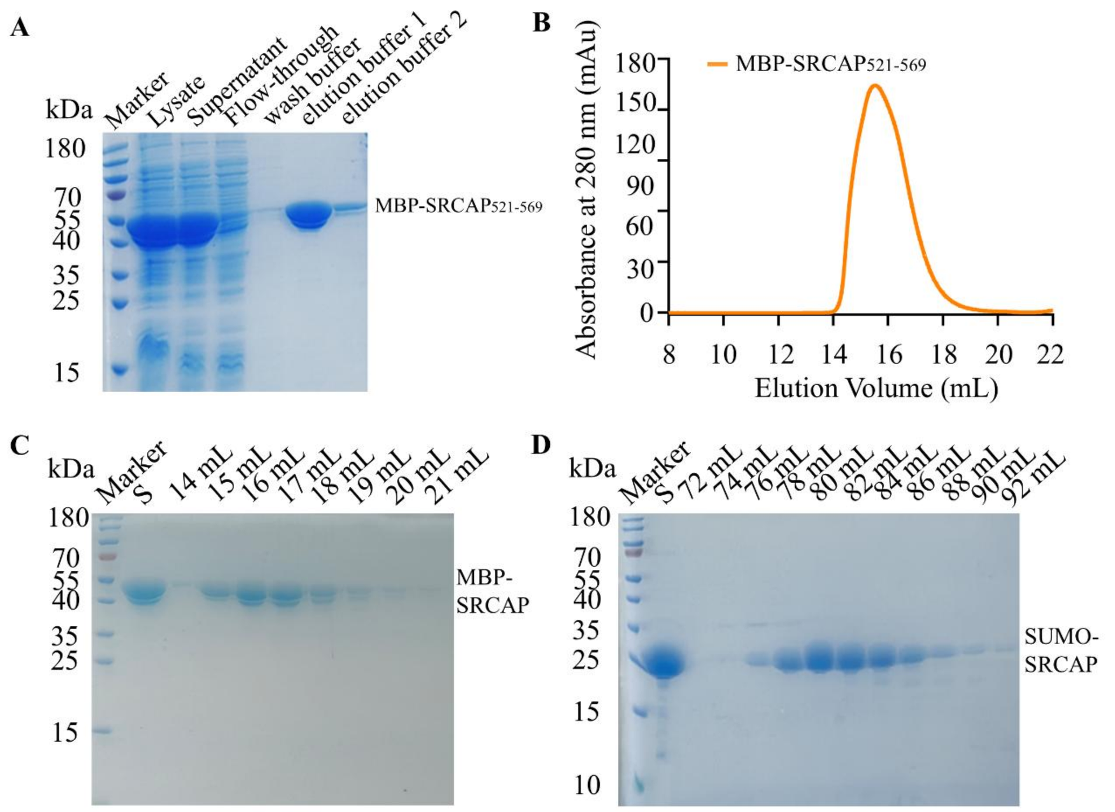


**Figure S2. Preparation of MBP-SRCAP$_{521-569}$ and SUMO-SRCAP$_{521-569}$.**

A) Purification of MBP-SRCAP$_{521-569}$ by Ni-NTA. Marker, protein standard molecular weight; Lysate, solution after centrifugation of lysed cells; Supernatant, Supernatant after centrifugation of lysed cells; Flow-through, supernatant liquid and unbound Ni beads; wash buffer, 25 mmol/L imidazole, 0.5 mol/L NaCl, 20 mmol/L Tris-HCl pH 8.0, 0.2% Glycerol, 1 mmol/L β-ME, 0.1 mol/L Urea; elution buffer 1, 250 mmol/L imidazole, 0.5 mol/L NaCl, 20 mmol/L Tris-HCl pH 8.0, 0.2% Glycerol, 1 mmol/L β-ME, 0.1 mol/L Urea; elution buffer 2, 500 mmol/L imidazole, 0.5 mol/L NaCl, 20 mmol/L Tris-HCl pH 8.0, 0.2% Glycerol, 1 mmol/L β-ME, 0.1 mol/L Urea;

B) Purification of MBP-SRCAP$_{521-569}$ by Superdex™ 200 Increase 10/300 GL.

C) MBP-SRCAP$_{521-569}$ analyzed by SDS-PAGE after gel filtration chromatography. Marker, protein standard molecular weight; S, sample prior to gel filtration chromatography; 14-21 mL, elution volume.

D) SUMO-SRCAP$_{521-569}$ analyzed by SDS-PAGE after gel filtration chromatography (HiLoad™ 16/600 Superdex™ 200 pg). Marker, protein standard molecular weight; S, sample prior to gel filtration chromatography; 72-92 mL, elution volume.

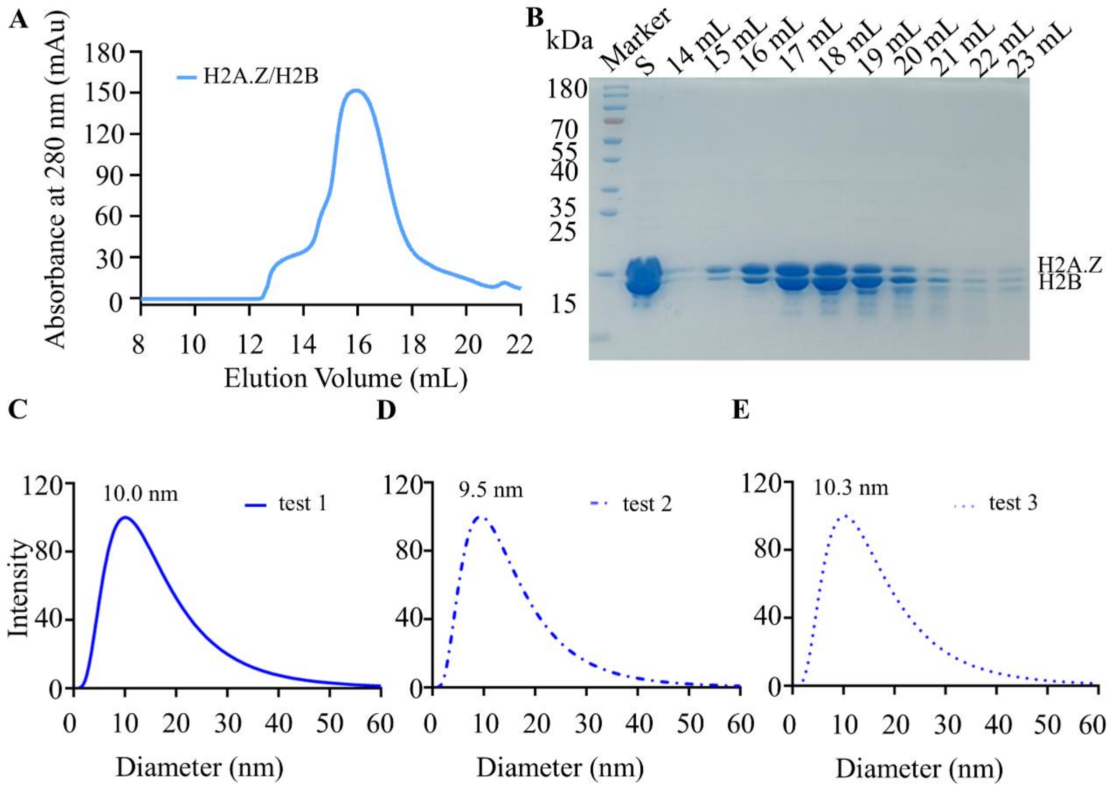


**Figure S3. Preparation of H2A.Z/H2B dimer and dynamic light scattering (DLS) analysis of the MBP-SRCAP$_{521\text{-}569}$/H2A.Z/H2B complex in solution.**

A) Purification of H2A.Z/H2B dimer by Superdex$^{TM}$ 200 Increase 10/300 GL.

B) H2A.Z/H2B dimer analyzed by SDS-PAGE after gel filtration chromatography. Marker, protein standard molecular weight; S, sample prior to gel filtration chromatography; 14-23 mL, elution volume.

C) DLS analysis 1 of the MBP-SRCAP$_{521\text{-}569}$/H2A.Z/H2B complex.

D) DLS analysis 2 of the MBP-SRCAP$_{521\text{-}569}$/H2A.Z/H2B complex.

E) DLS analysis 3 of the MBP-SRCAP$_{521\text{-}569}$/H2A.Z/H2B complex.

A

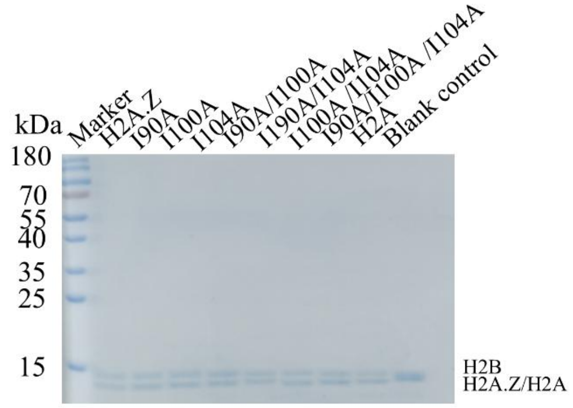


B

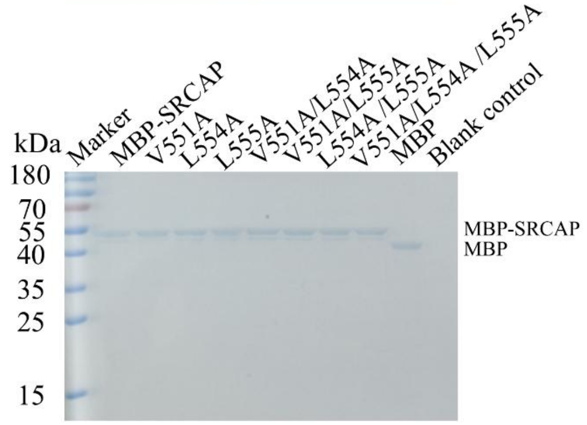


**Figure S4. Input of H2A.Z/H2B (H2A.Z WT or mutants) dimer or MBP-SRCAP$_{521\text{-}569}$ (WT or mutants) for MBP-pulldown experiments.**

A) SDS-PAGE of the input sample. Marker, protein standard molecular weight; H2A.Z, wild-type H2A.Z/H2B dimer; single mutant (I90A, I100A, or I104A), double mutant (I90A/I100A, I90A/I104A, or I100A/I104A), and triple mutant I90A/I100A/I104A of H2A.Z/H2B dimer; H2A, wild-type H2A/H2B dimer; Blank control, without H2A.Z/H2B dimer.

B) SDS-PAGE of MBP-SRCAP. Marker, protein standard molecular weight; MBP-SRCAP, wild-type MBP-SRCAP; single mutant (V551A, L554A, or L555A), double mutant (V551A/L554A, V551A/L555A, or L554A/L555A), triple mutant V551A/L554A/L555A of MBP-SRCAP; MBP, wild-type MBP; Blank control, without MBP-SRCAP.

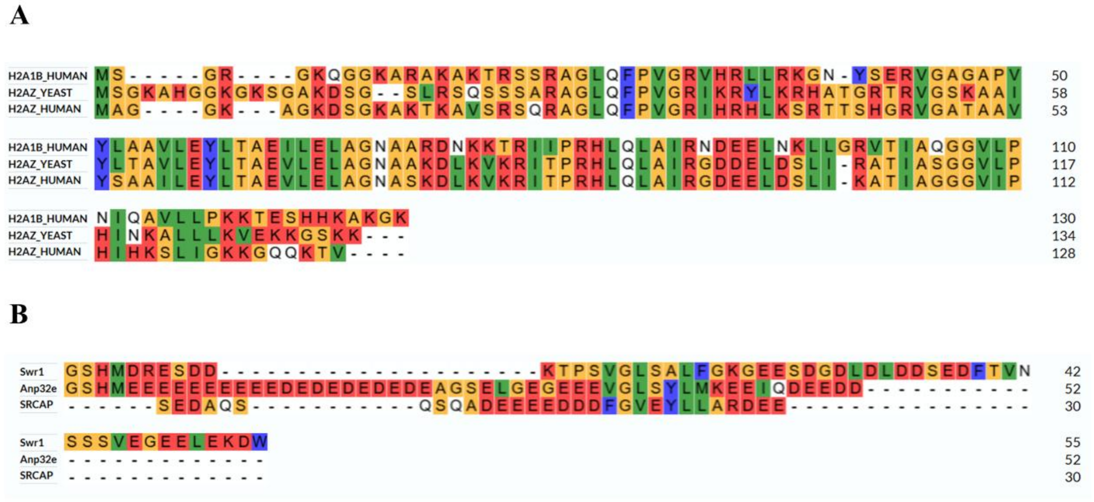


**Figure S5. Amino acid sequence alignment.**

A) Sequence alignment of full-length human H2A, yeast H2A.Z, and human H2A.Z.

B) Sequence alignment of the H2A.Z interacting domain (Z-domain) from yeast Swr1, human Anp32e, and human SRCAP.

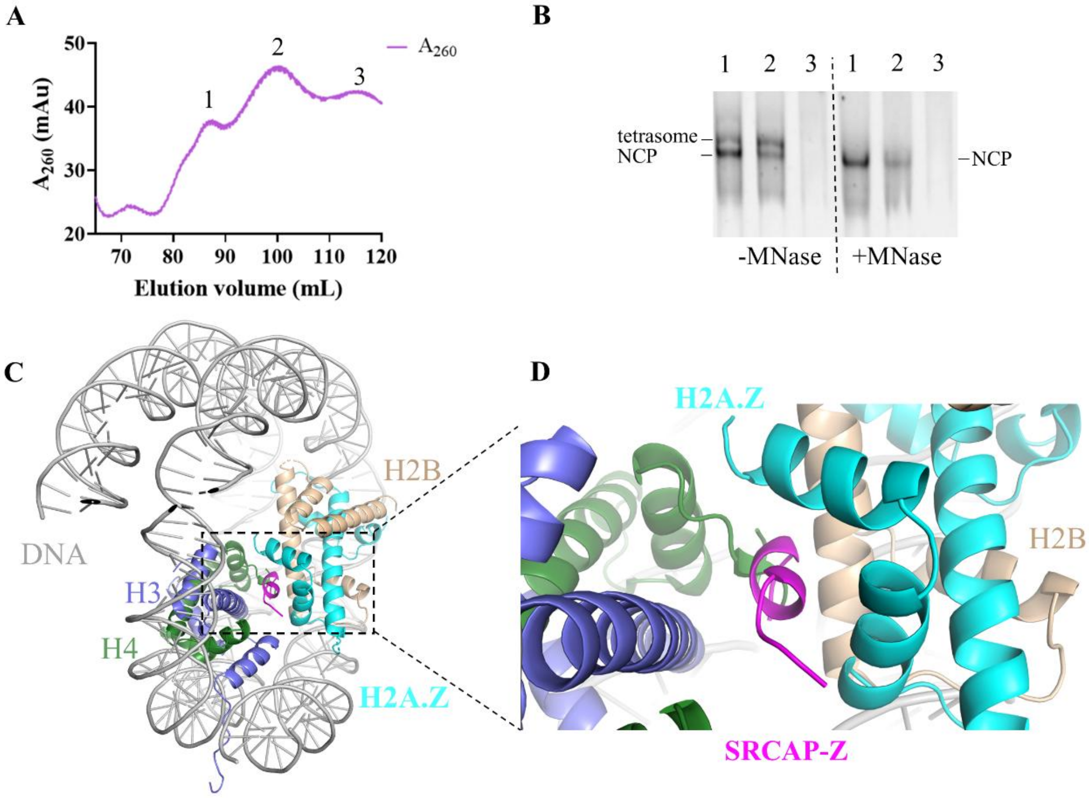


**Figure S6. Chaperone properties of the SRCAP-Z domain**

A) The DEAE exchange chromatography of nucleosome assembly of DNA-(H3.1-H4$^{A15C}$)$_2$ tetrasome and MBP-SRCAP$_{521-569}$/H2A.Z/H2B trimer.

SRCAP-Z promotes nucleosome formation. 1, main peak of nucleosome (NCP); 2, main peak of tetrasome; 3, DNA.

B) Collected samples at different elution positions, with or without digestion with 1.5 U MNase, and performing native PAGE. After digestion with MNase enzyme, the tetrasome band was gone (disappeared) because it was more loose than NCP, while the NCP band remained, indicating that the NCP is more resistant to MNase mediated digestion.

C) Overlay of the SRCAP-Z/scH2B-H2A.Z structure with the H2A.Z nucleosome structure (PDB ID: 1F66).

D) Enlarged details of SRCAP-Z interaction with histones in nucleosomes. SRCAP-Z may disturb the H3 (purple)-H4 (green) interaction with H2A.Z-H2B.

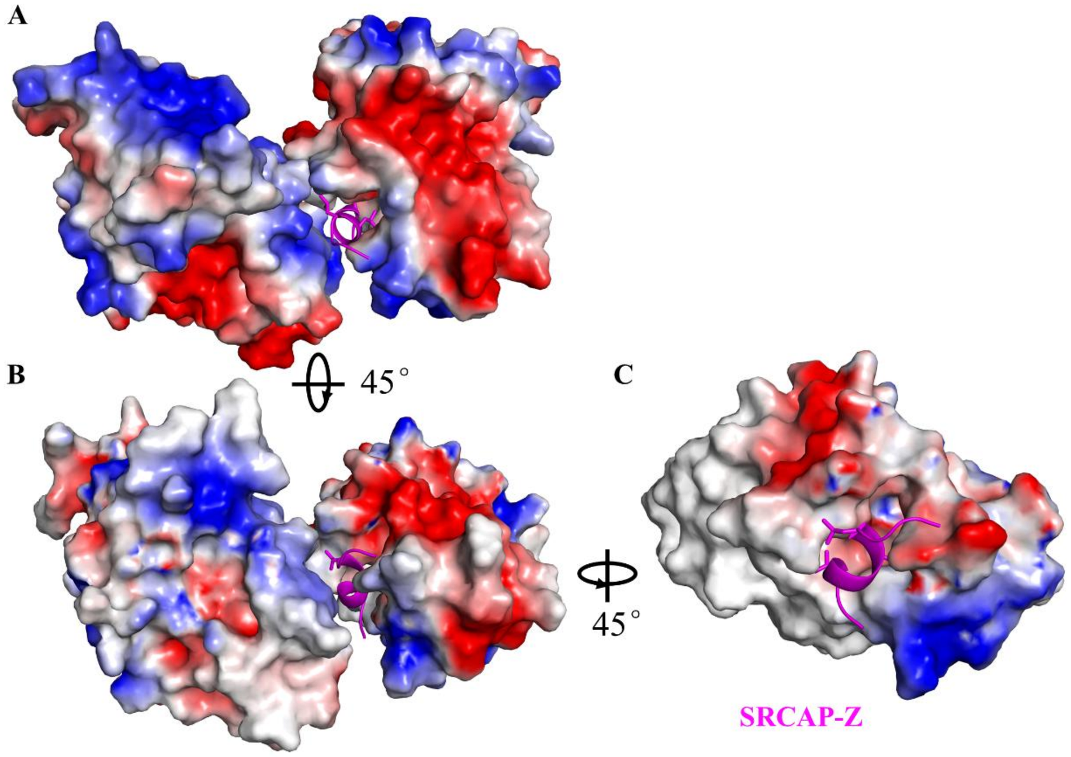


**Figure S7. The electrostatic distribution of the SRCAP-Z/scH2B-H2A.Z complex.**

A-C) Structure of the SRCAP-Z/scH2B-H2A.Z complex shown in electrostatic surface for histones at different angles. Red: negative charges, blue: positive charges.

**Table S1. Primers used for mutation and recombinant plasmid construction of $SRCAP_{521-569}$.**

Mutations in MBP-$SRCAP_{521-569}$ or SUMO-$SRCAP_{521-569}$ (Table S1).

Note:"F" denotes the forward primer, and "R" denotes the reverse primer. Mutation site dropdown line labeling.

**Table S2. Primers used for mutation of human H2A.Z.**

Mutations in H2A.Z (Table S2).

Note:"F" denotes the forward primer, and "R" denotes the reverse primer. Mutation site dropdown line labeling.

### EXPERIMENTAL PROCEDURES

#### Preparation of the $SRCAP_{531-560}$/scH2B-H2A.Z complex

The recombinant plasmid for scH2B-H2A.Z, stored in the laboratory, was transformed into *E. coli* BL21(DE3). Expression and purification were carried out according to previously described methods(Hong et al., 2014). Following quantification, the purified protein was mixed with $SRCAP_{531-560}$ (Peptide $SRCAP_{531-560}$ was synthesized by GL Biochem (Shanghai) Ltd.), concentrated, and stored frozen for future use.

#### Crystallization, diffraction data collection, and structure determination

The $SRCAP_{531-560}$/scH2B-H2A.Z protein was crystallized at 16°C using the vapor diffusion method. Crystals were initially screened in 96 well sitting drop plates and were obtained under a condition containing 0.2 mol/L potassium thiocyanate and 20% w/v polyethylene glycol 3350 (pH 7.0). Through optimization of the polyethylene glycol 3350 concentration, crystals suitable for data collection were obtained. The crystals were harvested and flash-frozen in liquid nitrogen (Supplementary Figure S1A), and subsequently shipped to the Shanghai Synchrotron Radiation Facility (SSRF) for X-ray diffraction data collection and analysis of the raw diffraction data

(Supplementary Figure S1B-C).

#### Cloning, expression, and purification of SRCAP recombinant protein

$SRCAP_{521-569}$ was amplified from a plasmid containing the SRCAP coding sequence and cloned into pET vectors containing MBP-tag or sumo-tag. Site-directed mutagenesis was performed to obtain mutant plasmids. The recombinant plasmids were transformed into the *E. coli* BL21(DE3) for low temperature soluble expression. At 18°C, expression was induced with a final concentration of 0.25 mmol/L IPTG for 16-18 hours. After induction, cells were harvested using a solution containing 25 mmol/L imidazole, 0.5 mol/L NaCl, and 20 mmol/L Tris-HCl (pH 8.0). Cells were lysed by sonication, and the supernatant of the lysate was purified by Ni-NTA (Qiagen) affinity chromatography (Supplementary Figure S2A), followed by further purification using gel filtration (HiLoad™ 16/600 Superdex™ 200 pg) (GE Healthcare) (Supplementary Figure S2B). Recombinant proteins were analyzed by SDS-PAGE (Supplementary Figure S2C-D).

#### Histone expression, purification, and assembly

The recombinant plasmids for H2A, H2A.Z, H2B, H3.1, and $H4^{A15C}$, stored in the laboratory, were transformed into *E. coli* BL21(DE3), respectively. Inclusion body forms of the histones were obtained by induction with 0.5 mmol/L IPTG at 37°C for various durations. Purification and assembly were performed according to our previously published article(Chittori et al., 2018) and our standard protocol of histone purification and mononucleosome assembly(Zhao and Hong, 2025) (Supplementary Figure S3).

#### Gel filtration chromatography analysis

The H2A.Z-H2B dimer and $MBP\text{-}SRCAP_{521-569}$ were mixed at a 1:1 molar ratio, and solid guanidine hydrochloride was added to a final concentration of 6 mol/L, followed by shaking to fully denature the proteins. Refolding was performed by dialysis to refolding buffer (0.2 mol/L NaCl, 10 mmol/L PBS pH 7.2, 1 mmol/L EDTA) at 4°C.

The refolded samples were subsequently analyzed by gel filtration chromatography. H2A.Z-H2B (Supplementary Figure S3) and MBP-SRCAP$_{521-569}$ were subjected to gel filtration chromatography analysis, respectively.

**MBP-pulldown Assay**

The interaction between MBP-SRCAP$_{521-569}$ and H2A.Z/H2B was examined *in vitro* by exploiting the specific binding affinity of maltose-binding protein (MBP) to amylose resin (Amylose Resin). MBP-tagged proteins were bound to the resin in a buffer containing 0.3 mol/L NaCl, 10 mmol/L PBS (pH 7.2), and 1 mM EDTA, followed by incubation at 4°C for 2 hours. After binding, the resin was washed with buffer to remove unbound proteins. Histones without the MBP tag were then added for binding. Following binding, the resin was washed with buffer. Finally, the bound complexes were eluted using an elution buffer (0.2 mol/L NaCl, 7 mol/L urea, 10 mmol/L maltose) with incubation in a 37°C water bath for 10 minutes. Blank controls consisting of MBP protein alone and the H2A/H2B dimer were included during the assay. The input/eluted samples were analyzed by SDS-PAGE (Supplementary Figure S4).

**Isothermal Titration Calorimetry (ITC) experiment**

The experiment was conducted using a MicroCal™ ITC200 isothermal titration calorimeter. Both the SUMO-SRCAP$_{521-569}$ and the H2A.Z/H2B dimer were dialyzed into the same buffer (0.2 mol/L NaCl, 10 mmol/L PBS pH 7.2). The reaction temperature was set up at 20°C, and the stirring speed was set up at 750 rpm. The instrument automatically titrated the SUMO-SRCAP$_{521-569}$ into the H2A.Z/H2B dimer solution and recorded the heat changes in real time. The data were analyzed using ITC200 analyze software according to the standard protocol of manufacture (https://www.malvernpanalytical.com.cn/).

**SRCAP function assay**

The H2A.Z/H2B dimer added MBP-SRCAP$_{521-569}$, was incorporated into DNA-(H3.1-H4$^{A15C}$)$_2$ tetrasome *in vitro*. The nucleosome core particles (NCP), tetrasomes, and free DNA were separated by DEAE ion-exchange chromatography(Hong et al., 2014) (Supplementary Figure S6). The isolated fractions were then subjected to electrophoretic mobility shift assay (EMSA) combined with micrococcal nuclease (MNase) digestion experiments to qualitatively assess the *in vitro* nucleosome assembly activity(Hong et al., 2014 , Yuan and Hong, 2025).
